# ZmEREB57 regulates OPDA synthesis and enhances salt stress tolerance through two distinct signal pathways in *Zea mays*

**DOI:** 10.1101/2022.07.05.498890

**Authors:** Jiantang Zhu, Xuening Wei, Chaoshu Yin, Hui Zhou, Jiahui Yan, Wenxing He, Jianbing Yan, Hui Li

## Abstract

In plants, APETALA2/Ethylene-Responsive Factor (AP2/ERF)-domain transcription factors play important roles in regulating responses to biotic and abiotic types of stress. In this study, a novel AP2/ERF gene in maize, ethylene responsive element binding factor 57 (*ZmEREB57*), was identified by comparative transcriptomics, which was significantly upregulated under salt treatment; however, its function remained unclear. Our findings revealed that ZmEREB57 is a nuclear protein with transactivation activity, induced by a number of abiotic types of stress. What’s more, two CRISPR/Cas9 knockout lines of *ZmEREB57* conferred sensitivity to saline conditions. DNA affinity purification sequencing (DAP-Seq) analysis revealed that ZmEREB57 mainly binds to promoters containing the O-box like motif (CCGGCC) to regulate the expression of its target genes. Further experiments validated that ZmEREB57 directly bind to the promoter of *ZmAOC2* involved in the synthesis of 12-oxo-phytodienoic acid (OPDA) and JA. Besides, transcriptome analysis identified a number of genes responded to both OPDA and JA treatment, while a different set of genes specifically responded specifically to OPDA, but not JA. Analysis of mutants deficient in the biosynthesis of OPDA and JA revealed that OPDA functions as a signaling molecule in the salt response. Our results provide evidence for ZmEREB57’s involvement in salt tolerance by regulating OPDA and JA signaling, and confirm early observations that OPDA signaling can function independently of JA signaling in maize.

## Introduction

Abiotic types of stress, such as high salinity, drought, and extreme temperatures, can adversely affect the growth and productivity of plants, particularly crops. Plants have thus developed a diverse set of mechanisms to cope with such inhibiting environmental conditions (Bartels & Sunkar, 2005). Among these, jasmonates (JAs) are lipid-derived signal transducer molecules known to have important roles in plant development and in mediating responses to biotic and abiotic stresses (Wasternack & Hause, 2013). JAs have been reported to accumulate in response to conditions such as drought and high salinity, leading to the expression of stress-response genes (Jung et al., 2007): the endogenous JA content in a salt-tolerant rice cultivar was found to be higher than in the salt-sensitive strain (Kang et al., 2005), and JA biosynthesis in rice roots has been reported to be strongly induced by drought stress (Takeuchi et al., 2011).

The only cyclopentenone precursor of JA is 12-oxo-phytodienoic acid (OPDA), which is synthesized via the α-linolenic acid metabolic pathway. Lipases aid the release of α-linolenic acid from plastidial membrane lipids, and this is oxygenated by 13-lipoxygenases (LOX) to form 13-hydroperoxylinolenic acid (Wasternack & Hause, 2013), which in turn generates 12, 13-epoxyoctadecatrienoic acid in the chloroplasts, catalyzed by allene oxide synthase (AOS). This is then used as a substrate by allene oxide cyclase (AOC) to form OPDA (Dave & Graham, 2012; Wasternack & Strand, 2016), which is transported into the peroxisome, where it undergoes reduction by OPDA reductase 3 (OPR3) and β-oxidation to generate JA (Dave & Graham, 2012; Kienow et al., 2008).

The bioactive form of JA - jasmonoyl-L-isoleucine (JA-Ile) - mediates the interaction between coronatine-insensitive protein 1 (COI1) with the JA-ZIM domain (JAZ) repressor to form the COI1-JAZ co-receptor complex (Staswick & Tiryaki, 2004). Under low JA conditions, JAZ proteins act as transcriptional repressors of gene expression. In response to elevated JA levels, the JA-Ile-mediated COI1-JAZ interaction triggers the ubiquitination of JAZ repressors and subsequent degradation by the proteasome, activating several transcription factors (TFs) that regulate specific physiological responses (Thines et al., 2007; Zhang et al., 2017).

Studies have suggested that OPDA is also involved in biotic and abiotic stress responses, embryo development, and seed germination (Satoh et al., 2014; Savchenko & Dehesh, 2014; Savchenko et al., 2014). However, several processes and genes are specifically activated by OPDA in a JA-Ile- and COI1-independent manner (Taki et al., 2005). Microarray analyses of Arabidopsis gene response to JA, methyl JA (MeJA), and OPDA treatments have revealed a set of genes that respond specifically to OPDA, but not to JAs (Stintzi et al., 2001). OPDA treatment of *coi1* mutants in Arabidopsis demonstrated that OPDA-specific gene expression is independent of the COI1-dependent JA signaling pathway (Taki et al., 2005). OPDA, but not JA, is found in *Marchantia polymorpha* and *Physcomitrella patens* exhibiting specific functions in defense and development (Yamamoto et al., 2015; Stumpe et al., 2010).

One of the largest groups of transcription factors (TFs) in plants is the APETALA2/Ethylene-Responsive Factor (AP2/ERF) superfamily. AP2/ERF TFs contain the highly conserved AP2 DNA-binding domain (∼60-amino acids), which binds the GCC-box (GCCGCC) and/or dehydration-responsive element (DRE)/C-repeat element to directly regulate downstream genes, serving important roles in plant development, defense against pathogens, and stress response (Sakuma et al., 2002; Mizoi et al., 2012; Xu et al., 2011). Certain ERF family members also respond to phytohormones and abiotic stress signals (Sugano et al., 2013). For example, rice *OsEBP2*, an ERF family member, was induced in response to compatible interaction with blast fungus and transiently induced by treatments with MeJA, abscisic acid (ABA) and ethylene, indicating *OsEBP2* possible role in abiotic stress (Lin et al., 2007). Furthermore, under conditions of wounding or water stress, the Arabidopsis octadecanoid-responsive AP2/ERF-domain TF 47 (AtORA47) is involved in the biosynthesis of JA, ABA and possibly ethylene, salicylic acid and strigolactone (Chen et al., 2016).

In the present study, the ethylene responsive element binding factor 57 (*ZmEREB57*) in maize was isolated and investigated in relation to response to salt stress, and two CRISPR/Cas9 knockout lines of *zmereb57* showed more sensitive to salt stress. We herein present evidence that ZmEREB57 binds the O-box like element in the promoter of *ZmAOC2*, regulating its expression and, in turn, the biosynthesis of OPDA and JA, which feed into two separate salt stress-response signaling pathways.

## Results

### Sequence analysis of ZmEREB57

RNA-seq studies was performed to determine the comparative transcriptomics under salt stress conditions (200 mM NaCl treatment) in maize inbred KN5585 line. In total, 12, 146 differentially expressed genes (DEGs) were identified (Supplementary Table S1). Among those DEGs, a gene *Zm00001eb193060*, encoding ethylene responsive element binding factor 57 (*ZmEREB57*) belonging to the AP2/ERF TF family, was significantly up-regulated under salt treatment (Fold change = 3.6356, *P* = 0.0002). To examine the role of the gene under salt stress condition, the full CDS of *ZmEREB57* was isolated basing on the transcriptomes of maize. The gene contained an open reading frame of 705 bp, encoding 234 amino acids, and the deduced protein sequence included the conserved AP2/ERF domain (Supplemental Figure S1a). The sequence alignment and phylogenetic analysis of ZmEREB57 and proteins from the AP2/ERF family from Arabidopsis, rice, soybean and Brachypodium indicated a high similarity among these homologs (Supplemental Figure S1a and b). Our analysis revealed 31.9% amino acid sequence identity between the AP2/ERF domain of ZmEREB57 and AtORA47 (the octadecanoid-responsive AP2/ERF-domain TF), as well as highly similar predicted three-dimensional structures, containing a long C-terminal α-helix and a three-stranded anti-parallel β-sheet (β1–β3) (Supplemental Figure S1c). These results indicate that ZmEREB57 is a member of the AP2/ERF group of TFs.

### ZmEREB57 localizes to the nucleus and exhibits transcription activation activity

To determine the subcellular localization of ZmEREB57, a ZmEREB57-GFP fusion protein expressed under the control of the CaMV *35S* promoter was transformed into tobacco epidermal cells. The microscopy images showed that the ZmEREB57-GFP construct induced fluorescence only in the nuclei, while the fluorescent signals of the control GFP construct were observed in the entire cells (Figure 1a). These indicated that the ZmEREB57 was localized at the nucleus.

**Figure 1.**
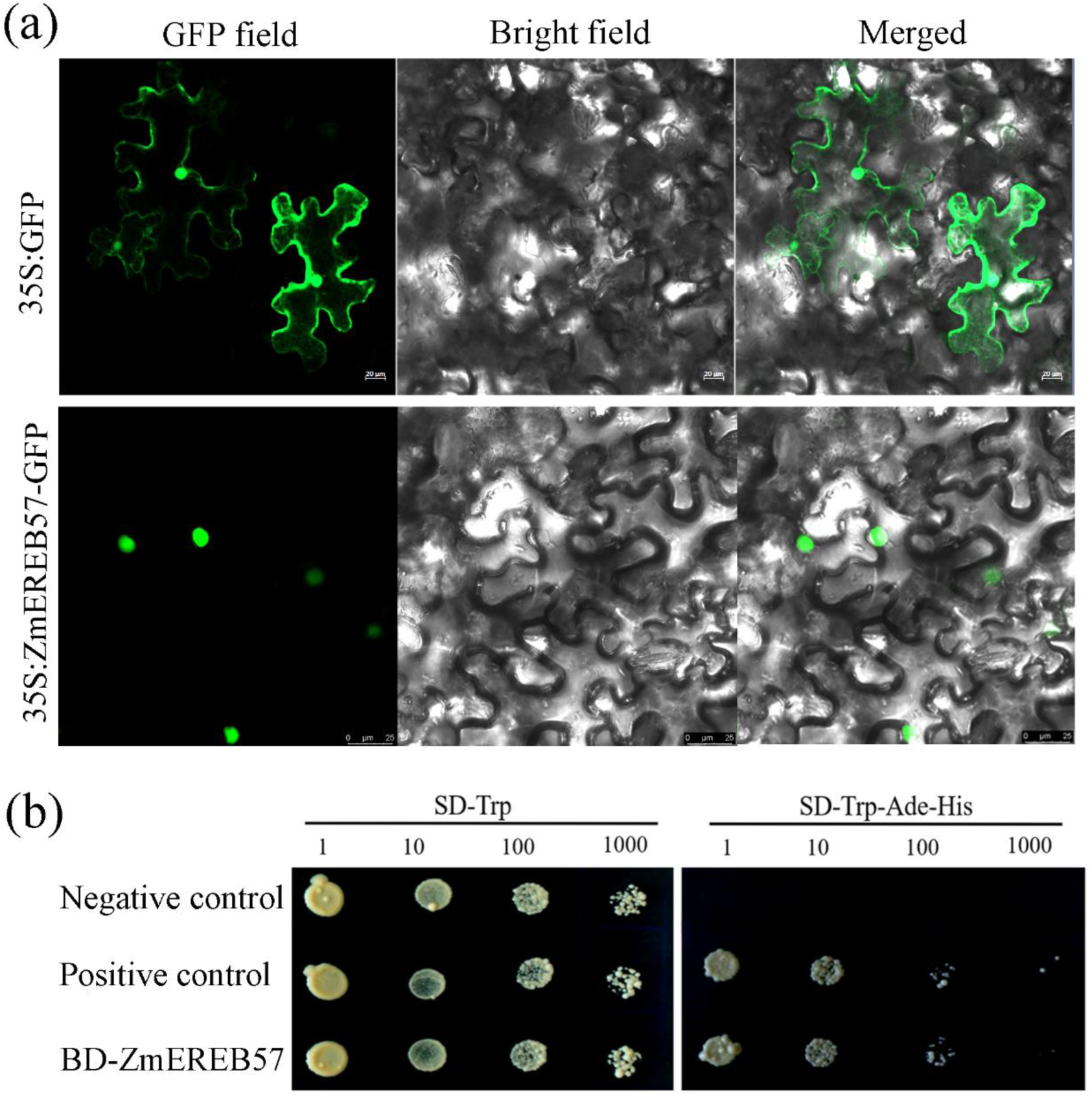
ZmEREB57 subcellular localization and transactivation activity assays. (a) The subcellular localization of ZmEREB57-GFP in *Nicotiana tabacum* epidermal cells was visualized using confocal laser scanning microscopy. (b) Transactivation assay of ZmEREB57. The construct of pGBKT7-ZmEREB57 was transformed into the yeast AH109 strain, which was cultured on SD/-Trp and SD/-Trp/-Leu/-Ade plates. GFP, green fluorescent protein; ZmEREB57, *Zea mays* Ethylene Responsive Element Binding Factor 57.

To determine whether ZmEREB57 acts as a transcription activator, a transactivation assay was performed in yeast (Figure 1b). All yeast cells containing pBD-ZmEREB57 recombinant construct, positive controls and negative controls could grow on the SD/-Trp medium (Figure 1b). By contrast, only transformants containing the pBD-ZmEREB57 vector and positive controls were able to grow on the more stringent media (SD/- Trp/-His/-Ade plates) (Figure 1b), indicating that ZmEREB57 was able to activate transcription in yeast.

### Expression of *ZmEREB57* can be induced by multiple types of abiotic stress

qPCR was performed to determine the *ZmEREB57* transcript levels in maize plants under different types of abiotic stress (Supplemental Figure S2). In response to drought and salt stress, a sharp increase in *ZmEREB57* transcription was observed 3 h after treatment, peaking at 12 h. Under oxidative stress, *ZmEREB57* transcripts of began to accumulate within 3 h and reached a maximum level at 24 h. The ABA and SA treatments resulted in similar effects, revealing a high level of *ZmEREB57* expression in response to these hormones, with transcription remaining higher than baseline even 24 h after phytohormone treatment application.

### Loss of *ZmEREB57* function renders maize plants sensitive to salt stress

Following the results that *ZmEREB57* expression was upregulated under conditions of high salinity, we generated two independent mutant lines using CRISPR-Cas9 system: *ZmEREB57^crispr^*-1 (*zmereb57-1*) and *ZmEREB57^crispr^*-2 (*zmereb57-2*), which conferred a 2- and 4-bp deletion, respectively, causing frameshifting and truncation of the resulting protein and the abundance of *ZmEREB57* transcript in mutants being significantly lower than in WT maize (Figure 2a and b; Supplemental Figure S3). We investigated the effects of salt stress on the *zmereb57* mutants grown in soil (Figure2c). When 20-day-old hydroponically grown maize plants were treated with 200 mM NaCl for 6 days, both two CRISPR/Cas9 knockout lines exhibited notably enhanced sensitivity to salt stress compared with the WT, with enhanced withering and bleaching in their leaves (Figure 2c). Correspondingly, the drop in relative water content and chlorophyll content of the two mutant lines was more notable than that in the WT (Figure 2d and e). These results demonstrate that the *zmereb57* mutant lines have decreased tolerance to salt stress than the WT.

**Figure 2.**
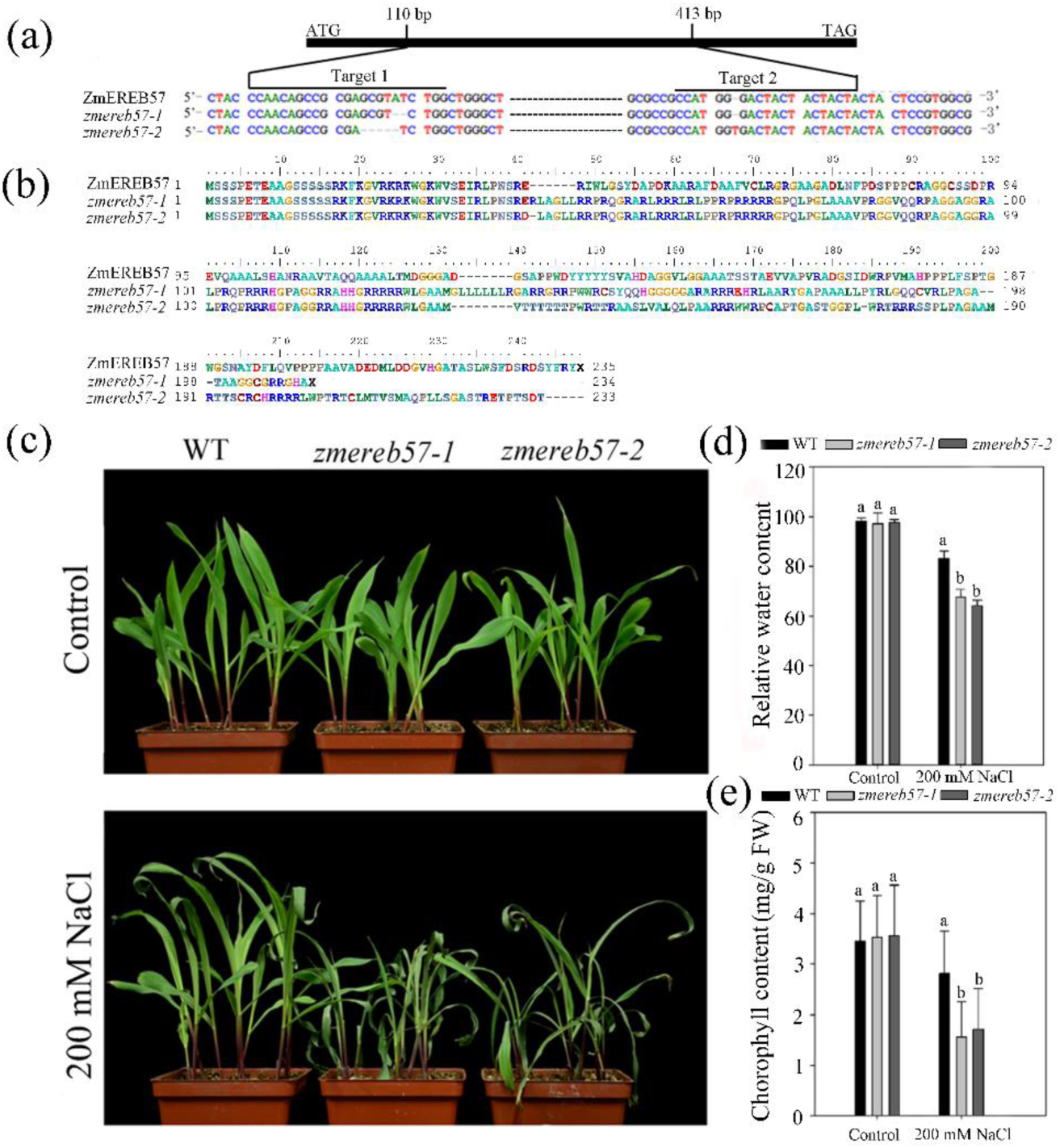
Analysis of *ZmEREB57* mutated lines. (a) Targeted mutagenesis of *ZmEREB57* via CRISPR-Cas9. Mutations in two independent lines (*zmereb57*-1 and *zmereb57*-2) are observed, following Sanger sequencing. (b) Alignment of the resulting amino acid sequences of ZmEREB57, *zmereb57*-1 and *zmereb57*-2. Only the sequences flanking the mutations are shown. (c) Appearance of 20 days-old mutated and WT plants grown in control and saline soil conditions (supplemented with 200 mM NaCl at day 6). The relative water content (d) and chlorophyll content (e) were determined in the WT and mutant plants following the salt treatments. Results are expressed as the mean ± SD (n=3 replicates). Different letters indicate values which departed significantly from those of WT maize. WT, wild type; ZmEREB57, *Zea mays* Ethylene Responsive Element Binding Factor 57.

To further investigate whether the expression of *ZmEREB57* occurs in response to salt stress, we analyzed the performance of Arabidopsis *ZmEREB57* overexpression lines under salt stress. In plants whether grown on MS medium or on pindstrup medium under control conditions, no phenotypic difference was observed between the transgene and WT lines. However, in plants grown on MS medium with 100 mM NaCl, both overexpression lines were able to develop larger leaves and longer roots, as well as retain more fresh weight than the WT (Supplemental Figure S4a–c). Upon treatment with 200 mM NaCl, most of the WT plants grown on pindstrup medium wilted and eventually perished at the vegetative growth stage. By contrast, the plants of the overexpression lines generally maintained a healthy green appearance and only slight wilting of the leaves was observed (Supplemental Figure S4d). Correspondingly, the fresh weight and chlorophyll content of the two overexpression lines under salt stress was significantly higher than those of the WT plants (Supplemental Figure S4e and f). Notably, complementation of the *atora47* mutant with overexpression of *ZmEREB57* appeared to fully restore salt tolerance to levels similar to those of the WT (Supplemental Figure S4a and d), including an increase in root length, fresh weight and chlorophyll content of the seedlings to WT levels under conditions of salt stress (Supplemental Figure S4b-f). These data imply that *ZmEREB57* overexpression enhances tolerability to salinity stress.

### Identification of ZmEREB57 target genes in maize

To investigate the target genes regulated by ZmEREB57, DAP-Seq was performed for this purpose. As a result, total 606 potential ZmEREB57 directly targeted genes with functional annotations were detected across the whole maize genome (Figure 3a). Among the 606 genes, approximately 18% (112) were located within the promoter region (2 kb upstream from the TSS) (Figure 3b, Supplemental Table S2). From promoter binding sites, a specific core binding motif CCGNCC showed the most significance (Figure 3c), which belonged to the ‘O-box like’ sequence. 112 promoter genes were mainly 7 transcription factors, 9 stress responsers, 28 enzymes involved in biological processes, 47 proteins involved in cellular component, and 21 uncharacterized proteins (Figure 3d). These results suggested that ZmEREB57 is a transcription factor with multiple regulatory responses.

**Figure 3.**
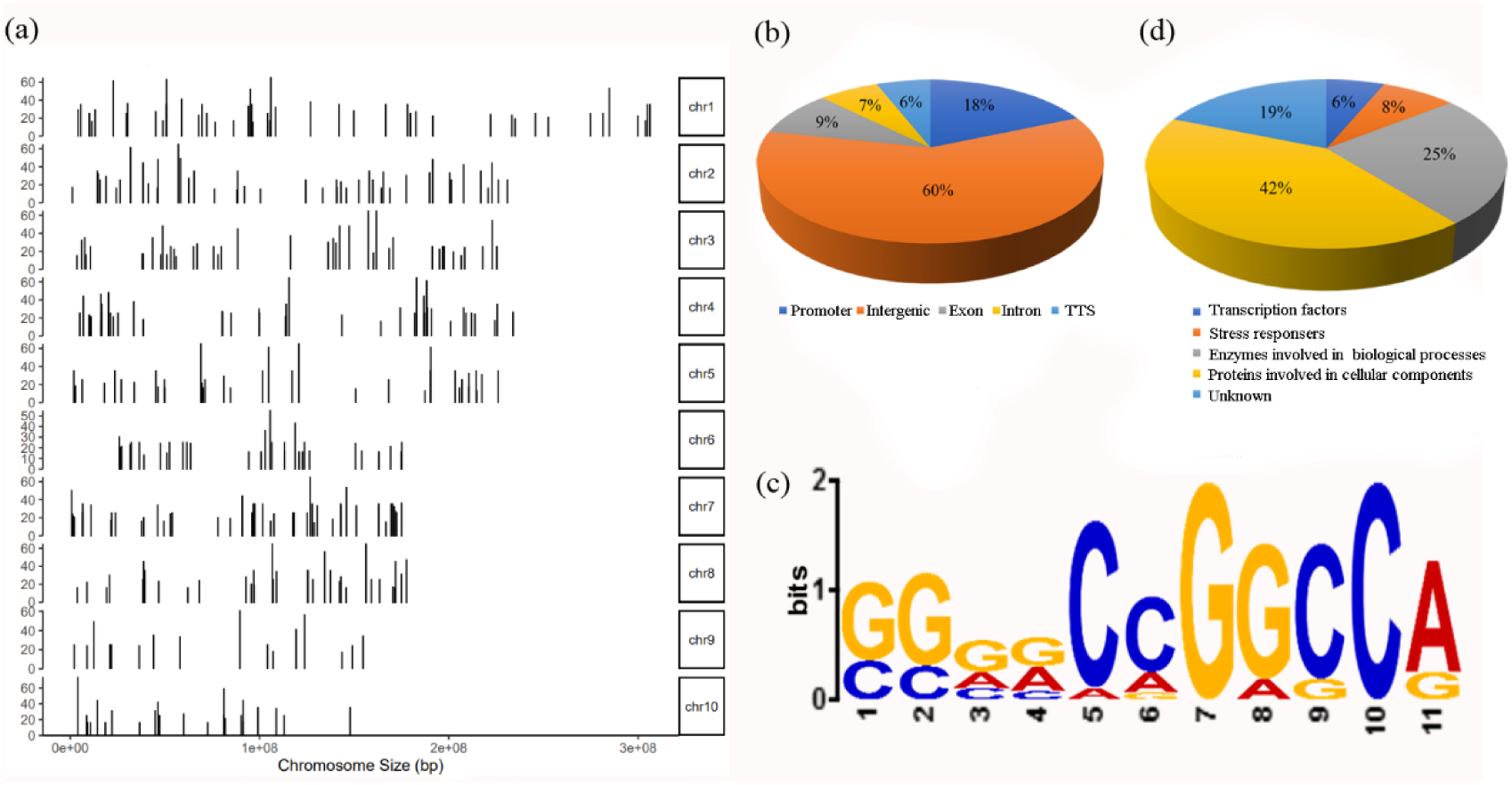
DAP-seq (DNA affinity purification sequencing) analysis of ZmEREB57 target genes. (a) Distribution of ZmEREB57-binding sites along the ten chromosomes of maize. (b) Distribution of ZmEREB57-binding sites in genic and intergenic regions. (c) Motif analysis of combined peaks of ZmEREB57 with the most significant *E*-value. (d) Biological process categorization of ZmEREB57-regulated target genes.

### ZmEREB57 activates *ZmAOC2* expression by directly binding to its promoter

Among the promoter genes bound by ZmEREB57, we found a gene that encoded allene oxide cyclase2 protein (*Zm00001eb393520*, ZmAOC2), involved in JA synthesis, was examined with a strong enriched peaks from DAP-seq analysis (Supplemental Table S1). To confirm this binding, the putative ZmEREB57-binding elements in the promoter of *ZmAOC2* was searched and screened by means of a Y1H assay. The results revealed that an ‘O-box’ like sequence CCGGCC exists in the promoter region of *ZmAOC2*, and that ZmEREB57 was able to specifically bind to this element in yeast (Figure 4a), suggesting that *ZmAOC2* is a target gene of ZmEREB57. Moreover, luciferase and EMSA assays were performed to confirm the direct binding of ZmEREB57 to the *ZmAOC2* promoter. The results of the luciferase assay showed that ZmEREB57 led to luciferase expression in the presence of the *ZmAOC2* promoter (Figure 4b). For the EMSA, a 40-bp oligonucleotide containing the CCGGCC sequence was synthesized based on the sequence of the *ZmAOC2* promoter, and was labeled as a probe. When the ZmEREB57 protein was incubated with the labeled probe, the migration speed of the protein-DNA complex was observed to be reduced (Figure 4c). Competition experiments were performed to determine the specificity of the mobility shift. When the ratio of unlabeled probe to labeled probe was 25:1 or 50:1, the majority of labeled probe was weakened (Figure 4c), indicating that the ZmEREB57 protein can bind specifically to the O-box-like in the *ZmAOC2* promoter. As expected, the luciferase assay and EMSA results revealed that ZmEREB57 can also bind to the *AtAOC2* promoter in vitro, which containing ‘O-box’ like sequence CCGGCC (Supplemental Figure S5).

**Figure 4.**
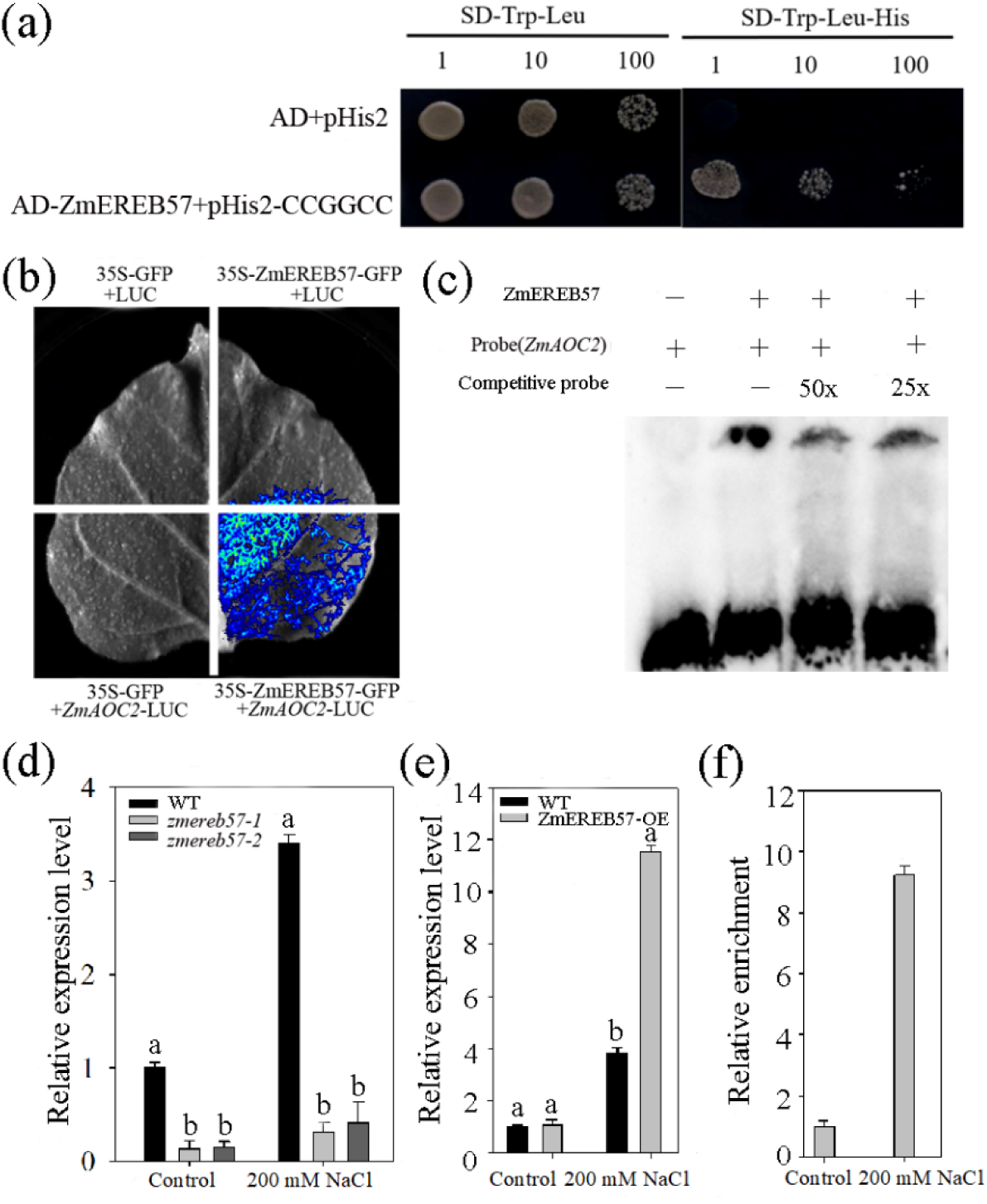
ZmEREB57 binds directly to the *ZmAOC2* promoter and activates its expression. (a) The binding activity of ZmEREB57 to the O-box sequence motif, as assessed using a yeast one-hybrid assay. Yeast cells were selected on SD/-Trp/-Leu and SD/-Trp/-Leu/-His media. (b) Luciferase assay for the *ZmAOC2* promoter. The pro*ZmAOC2*:LUC-35S:REN reporter construct was transiently expressed in tobacco leaves together with the control vector or the effector, and images of luciferase signal detection were captured. (c) An electrophoretic mobility shift assay indicates that ZmEREB57 binds to the *ZmAOC2* promoter *in vitro*. His-ZmEREB57 protein (2 μg) was incubated with labeled probe. Unlabeled (competitive) probe at 25x and 50x molar excess competed with the labeled probe. Relative expression levels of (d) *ZmAOC2* in WT and *zmereb57* mutant maize plants, and (e) *AtAOC2* in WT and transgenic Arabidopsis plants, with and without salt treatment. (f) Chromatin immunoprecipitation-quantitative PCR assay of chromatin isolated from the transgenic *ZmEREB57* overexpression line and immunoprecipitated with an anti-GFP antibody shows that ZmEREB57 binds to the *AtAOC2* promoter *in vivo*. Relative expression levels of *AtAOC2* were normalized against those in the no-antibody control. Results are expressed as the mean ± SD (n=3 replicates). Different letters indicate values which departed significantly from those of WT maize. AtAOC2, *Arabidopsis thaliana* allene oxide cyclase 2; WT, wild type; ZmEREB57, *Zea mays* Ethylene Responsive Element Binding Factor 57.

Further investigation included the analysis of *ZmAOC2* expression in WT and *zmereb57* CRISPR/Cas maize lines under control and salt stress conditions. Under control conditions, *ZmAOC2* was expressed at higher levels in the WT plants than in the *zmereb57* CRISPR/Cas lines (Figure 4d). Following salt treatment, the *ZmAOC2* expression level was further enhanced in the WT, while very little change was observed in the *zmereb57* mutants (Figure 4d). Accordingly, *ZmEREB57* overexpression in Arabidopsis resulted in enhanced *AtAOC2* expression under salt stress (Figure 4e). Finally, we performed ChIP-qPCR assays using the Arabidopsis *ZmEREB57* overexpression line under control and salt stress conditions. Under the control conditions, ZmEREB57 was found to bind the *AtAOC2* promoter, and salt treatment appeared to further enhance this binding (Figure 4f). Together, these data indicate that ZmEREB57 directly binds to the *ZmAOC2* promoter and acts as a transcription activator.

### ZmEREB57 affects the accumulation of endogenous OPDA and JA in maize

AOC has been reported as a dimeric enzyme localized in the chloroplasts, catalyzing the formation of OPDA from 13-hydroperoxide linolenic acid; OPDA is then converted into JA in the peroxisomes (Wu et al., 2011; Ziegler et al., 2000). To verify the involvement of *ZmAOC2* in the synthesis of OPDA and JA in maize, we first investigated the subcellular localization of ZmAOC2, where we observed the signal from the ZmAOC2-GFP fusion protein to be concentrated in the chloroplasts (Figure 5a). We then analyzed the OPDA and JA content in WT versus *zmaoc2* mutant plants under control and salt stress conditions. In the absence of salt stress, the OPDA and JA content in the *zmaoc2* mutant was lower than in the WT plants (Figure 5b and c), and treatment with 200 mM NaCl led to an even bigger difference between the OPDA and JA levels in the two lines, due to a marked increase in the content of these molecules in the WT (Figure 5b and c). These findings indicate that *ZmAOC2* is involved in the OPDA and JA synthesis in maize.

**Figure 5.**
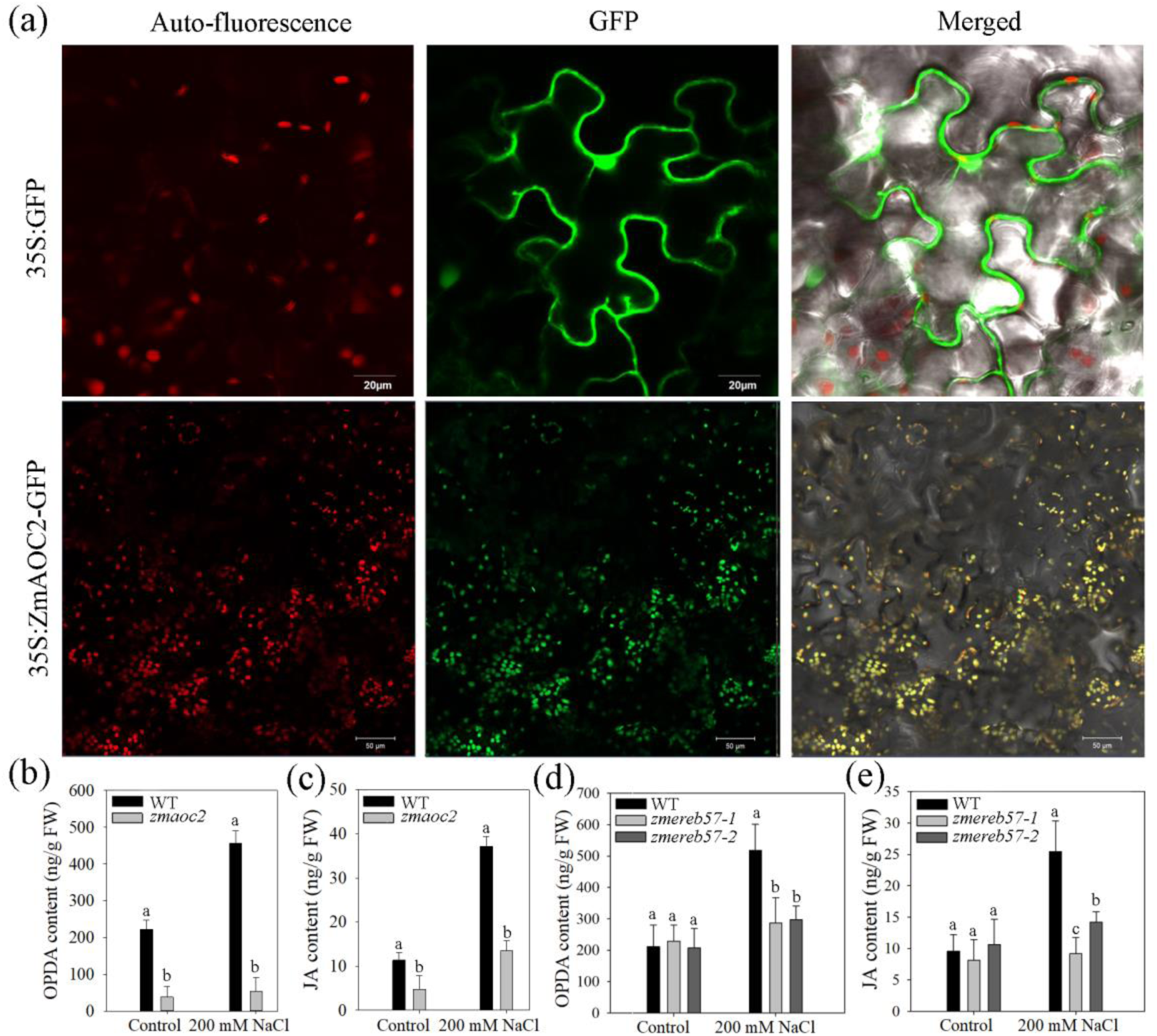
Effect of *ZmEREB57* expression on OPDA and JA levels. (a) The subcellular localization of ZmAOC2-GFP in *N. benthamiana* epidermal cells was visualized using confocal laser scanning microscopy, indicating localization in the chloroplasts. The levels of (b) OPDA and (c) JA in WT and a *zmaoc2* mutant maize line with or without 200 mM NaCl treatment. The levels of (d) OPDA and (e) JA in WT and *zmereb57* mutant maize lines with or without 200 mM NaCl treatment. Data are presented as the mean ± SD (n = 3 replicates). Different letters indicate values which departed significantly from those of WT maize. FW, fresh weight; GFP, green fluorescent protein; JA, jasmonate; OPDA, 12-oxo-phytodienoic acid; WT, wild type; ZmEREB57, *Zea mays* Ethylene Responsive Element Binding Factor 57.

Endogenous OPDA and JA levels were also quantified in WT and *zmereb57* mutant maize lines, where little difference was observed these plants under control conditions. By contrast, on application of salt stress, the OPDA and JA content increased more strongly in the WT than in the *zmereb57* mutants (Figure 5d and e). These results suggest that ZmEREB57 affects endogenous OPDA and JA accumulation by regulating *ZmAOC2* expression under conditions of salt stress.

### Exogenous OPDA or JA restore the salt-sensitive phenotype of *zmereb57* mutants

To demonstrate the effect of OPDA or JA on the sensitivity to salt stress, we applied exogenous OPDA or JA to WT and *zmereb57* mutant lines that were treated with 200 mM NaCl, and analyzed their phenotype. Under treatment with 200 mM NaCl without added OPDA or JA, *zmereb57* mutants exhibited severe wilting and higher water and chlorophyll loss compared with the WT plants (Supplemental Figure S6a–c). However, following application of 100 μm OPDA or 100 μm JA under conditions of salt stress, no phenotypic difference could be observed between the mutant lines and the WT (Figure 6a and f). The relative water content and chlorophyll levels decreased slightly after 48 h of NaCl treatment, but no significant difference was observed among the plant lines (Figure 6b, c, g, h). When plants were treated with NaCl as well as OPDA, the levels of endogenous OPDA and JA were both observed to increase in the WT and *zmereb57* mutant plants (Figure 6d and e). Notably, in plants treated with NaCl and JA, the levels of endogenous OPDA or JA increased equally in all strains, but endogenous OPDA levels increased only slightly in the *zmereb57* mutant plants compared with the WT (Figure 6i and j). These results indicate that exogenous OPDA or JA application could help restore the salt-tolerant phenotype of *zmereb57* mutants by leading to an increase in the levels of endogenous OPDA and JA.

**Figure 6.**
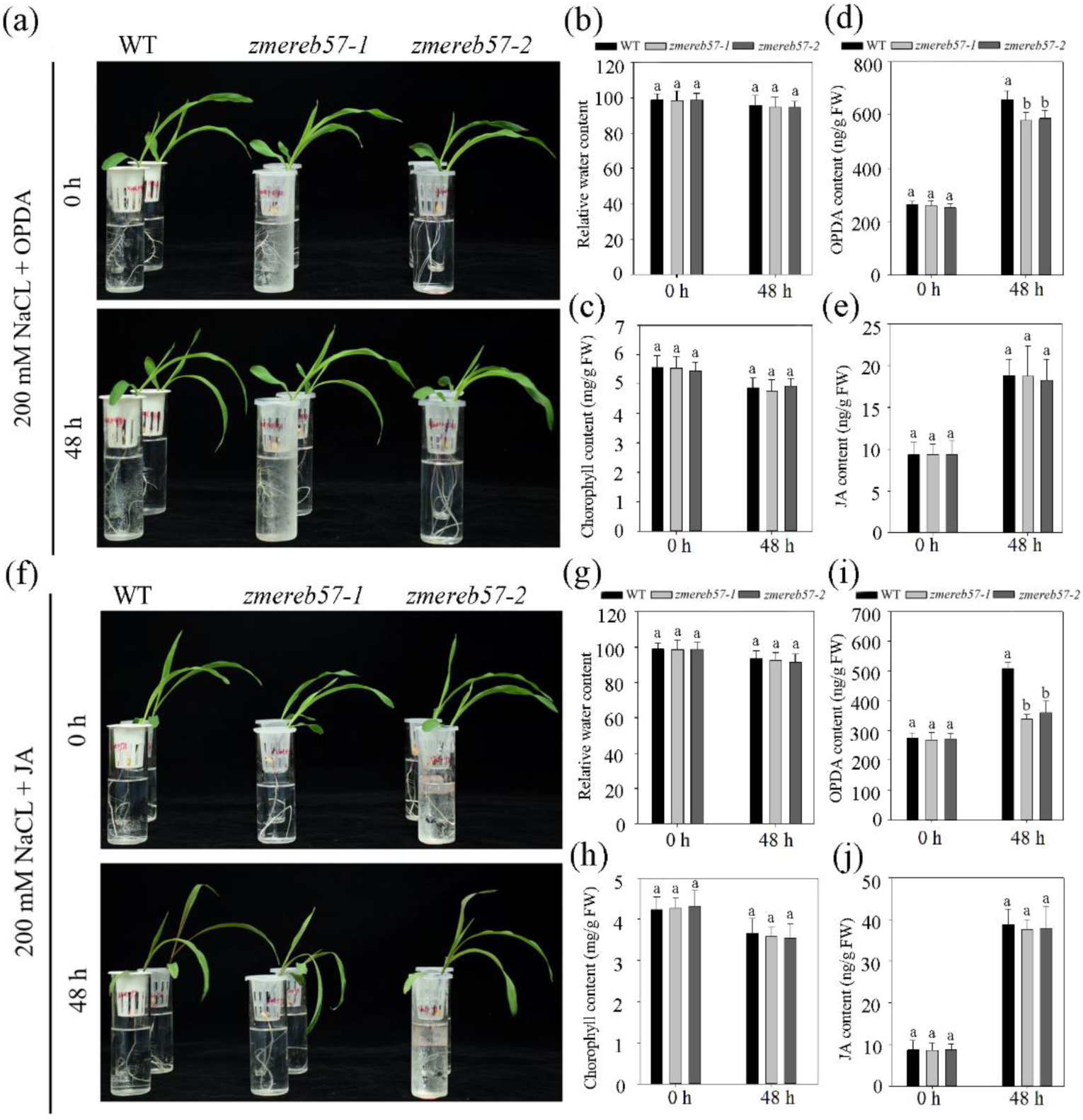
Effect of exogenous OPDA or JA on the sensitivity of *zmereb57* mutant plants to salt stress. (a) Appearance of WT and *zmereb57* mutant maize plants grown in high salt conditions and supplemented with 100 μM OPDA. Measurements of (b) relative water content, (c) chlorophyll, (d) OPDA, and (e) JA levels were obtained for WT and *zmereb57* mutant plants after salt with OPDA treatment. (f) Appearance of WT and *zmereb57* mutant maize plants grown in high salt conditions and supplemented with 100 μM JA. Measurements of (g) relative water content, (h) chlorophyll, (i) OPDA, and (j) JA levels were obtained for WT and *zmereb57* mutant plants after salt + JA treatment. Data are presented as the mean ± SD (n = 3 replicates). Different letters indicate values which departed significantly from those of WT maize. FW, fresh weight; JA, jasmonate; OPDA, 12-oxo-phytodienoic acid; WT, wild type; ZmEREB57, *Zea mays* Ethylene Responsive Element Binding Factor 57.

### Genes differentially expressed under OPDA and JA treatment participate in the response to salt stress

To study the gene expression patterns associated with *ZmEREB57*-related regulation of OPDA/JA synthesis in response to salt stress in maize, RNA-seq analyses of WT plants treated with or without 100 μM OPDA or 100 μM JA were carried out, and was characterized the comparative transcriptomics combined with previous 200 mM NaCl treatment. Compared with grown under control conditions, the transcriptome analysis of maize leaves from plants subjected to OPDA or JA treatments for 48 h identified 6,911 and 11,958 DEGs, respectively (Figure 7a and Supplemental Table S3-4). Overall, 4,032 OPDA-specific response genes (ORGs, 58.3% in OPDA-related DEGs) and 5,614 JA-specific response genes (JRGs, 46.9% in JA-related DEGs) were found to be genes that also responded to salt treatment (Figure 7b and Supplemental Table S5-6). A total of 1,351 DEGs (OPDA and NaCl-specific response genes, O&NRGs) were common between OPDA- and NaCl-related transcripts (but not JA treatment), and 2,933 DEGs (JA and NaCl-specific response genes, J&NRGs) were found to be a result of JA as well as NaCl treatment (but not OPDA treatment) (Figure 7b). Of the DEGs identified in this analysis, 2,681 transcripts (OPDA and JA and NaCl-specific response genes, O&J&NRGs) were identified in all three categories, i.e. differentially expressed due to salt stress as well as OPDA and JA treatment (Figure 7b and Supplemental Table S7). These findings suggest that a considerable proportion of the maize transcriptome changes triggered by OPDA and/or JA treatment are involved in the plant’s response to salt stress.

**Figure 7.**
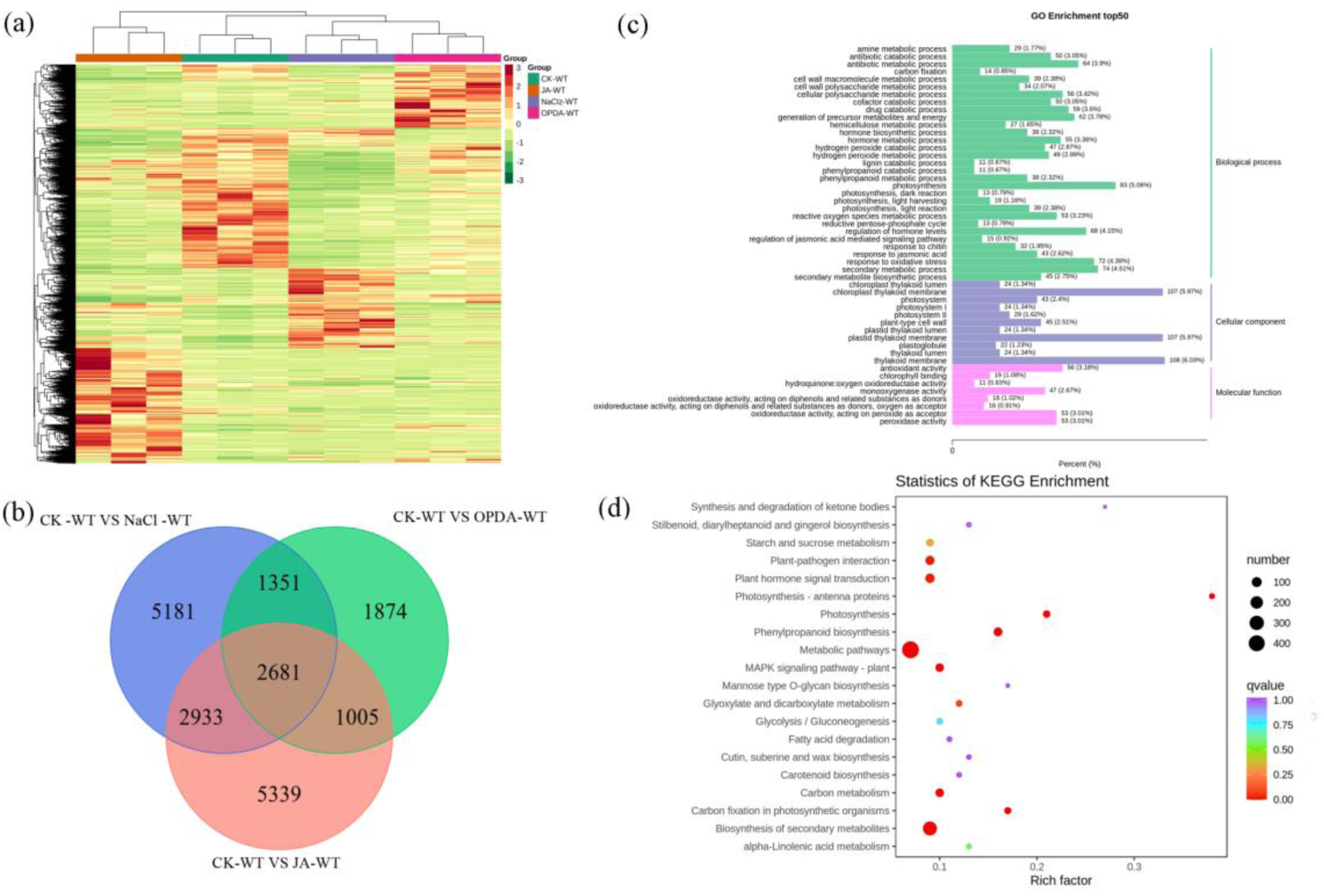
Global transcriptome response of maize to salt and phytohormone treatments. (a) DEGs identified in plants treated with 200 mM NaCl with or without 100 μM OPDA or 100 μM JA supplement. (b) Venn diagrams showing the overlap of DEGs identified from the different treatment conditions. (c) Gene ontology term enrichment annotation and (d) Kyoto Encyclopedia of Genes and Genomes pathway enrichment annotation of the 1,351 DEGs shared between treatments with 200 mM NaCl alone and 200 mM NaCl supplemented with OPDA, but not the 200 mM NaCl plus JA treatment. Numbers next to each bar represent the number of DEGs. DEG, differentially expressed gene; JA, jasmonate; OPDA, 12-oxo-phytodienoic acid.

To gain insight into the biological processes triggered in response to salt stress, we performed a GO term enrichment analysis of the 2,681 O&J&NRGs and the 1,351 O&NRGs (Figure 7c and d; Supplemental Figure S7). These two groups of DEGs included enrichment of genes involved in different biological processes, including defense response, cell wall organization, lipid catabolic processes, photosynthesis and salt stress response, as well as genes categorized with hormone-related GO terms, such as response to ABA and salicylic acid, or the ABA-mediated signaling pathway. Among the enriched GO terms of DEGs shared among all three treatments, the majority described genes encoding TFs. These TFs appeared to belong to several different families, including WRKY, myeloblastosis (MYB), NAC, AP2/EREBP, basic region leucine zipper (bZIP), basic helix-loop-helix, and MYC, which serve critical roles in plant responses to abiotic types of stress. Additionally, many genes related oxidative stress from DEGs were identified, for example, mitogen-activated protein kinase (Figure 7c and d; Supplemental Figure S7).

To confirm the robustness of the differential gene expression analysis, six O&J&NRGs associated with salt stress response processes were selected for reverse transcription-qPCR analysis in maize WT and *zmereb57* mutants, and the results were compared with the RNA-seq data. The expression of stress/ABA-response genes *RD29A* (*Zm00001eb339990*), *RD29B* (*Zm00001eb408850*), *ABF3* (*Zm00001eb032760*), *ABA1* (*Zm00001eb390480*), *DREB2A* (*Zm00001eb336270*) and *NCED3* (*Zm00001eb031940*) was strongly upregulated in the WT plants treated with 200 mM NaCl, but remained almost unaffected by the treatment in the *zmereb57* mutants under the same conditions (Supplemental Figure S8). The relative expression profiles of each gene were consistent with the differential expression detected by means of RNA-seq, supporting the accuracy of the transcriptome analysis results.

### OPDA enhances salinity tolerance through JA-dependent and -independent pathways

Due to the similarity in chemical structure between OPDA and JA, we hypothesized that OPDA-related responses may also be regulated by the JA-COI1 pathway in maize. To investigate whether OPDA confers tolerance to salt stress independently as well as cooperatively with JA, we utilized two maize mutants of JA biosynthesis: *aoc2*, which is unable to synthesize neither OPDA nor JA, and *opr3*, which contains a defect in JA biosynthesis. The effect of applying exogenous OPDA or JA on the *zmaoc2* or *zmopr3* mutants’ resistance to salt stress was tested. When the plants were treated with 200 mM NaCl alone, both the *zmaoc2* and *zmopr3* mutant strains exhibited more severe wilting and higher water and chlorophyll loss than the WT (Figure 8a–c). On the addition of 100 μm OPDA to this treatment, the characteristics of the mutant plants and the WT were similar (Figure 8d–f). However, when supplemented with 100 μM JA, only the *zmopr3* mutant exhibited an appearance and measurements similar to the WT phenotype, while the *zmaoc2* mutant appeared more sensitive (Figure 8g–i). These results indicate that OPDA enhances tolerance to salinity in pathways that are dependent and independent of JA.

**Figure 8.**
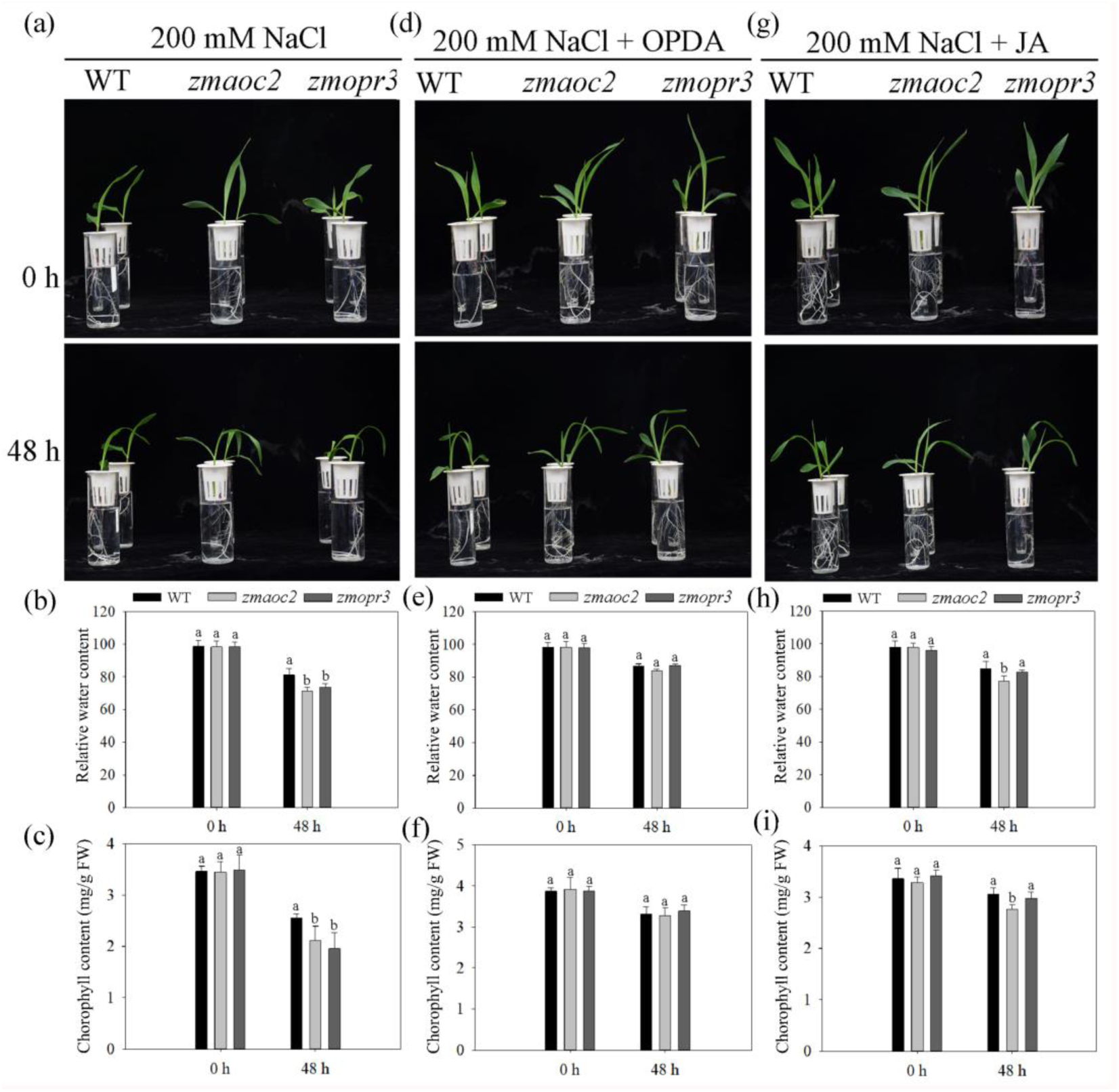
Characteristics of OPDA and JA biosynthesis mutants under conditions of salt stress. (a) Phenotype, (b) relative water content, and (c) chlorophyll levels of maize WT, *zmaoc2* and *zmopr3* mutants before and after treatment with 200 mM NaCl. (d) Phenotype, (e) relative water content, and (f) chlorophyll levels of maize WT, *zmaoc2* and *zmopr3* mutants before and after treatment with 200 mM NaCl supplemented with 100 μM OPDA. (g) Phenotype, (h) relative water content, and (i) chlorophyll levels of maize WT, *zmaoc2* and *zmopr3* mutants before and after treatment with 200 mM NaCl supplemented with 100 μM JA. Data are represented as the mean ± SD (n = 3 replicates). Different letters indicate values which departed significantly from those of WT maize. JA, jasmonate; OPDA, 12-oxo-phytodienoic acid; WT, wild type.

To determine whether OPDA is required to induce the expression of ORGs under conditions of salt stress *in vivo*, we assessed the expression profile of the selected five putative O&NRGs genes, including *ZAT10* (*Zm00001eb064620*), *FAD* (*Zm00001eb414630*), *ERF9* (*Zm00001eb369560*), *DREB1* (*Zm00001eb073550*), *GST6* (*Zm00001eb402640*), and one J&NRGs gene *VSP9a* (*Zm00001eb398120*) in the two mutant strains (Supplemental Figure S9). Under control conditions, the genes’ expression did not differ from that in the WT plants. Following salt treatment, the transcription levels of all genes were lower in the *zmaoc2* mutant plants compared with those seen in the WT. Supplementing the salt treatment with exogenous OPDA, but not JA, resulted in an increase of *ZAT10*, *FAD-OXR*, *ERF5*, *DREB2A* and *GST6* expression in both the WT and *zmaoc2* mutant strain. Expression of *VSP2* was induced by application of exogenous OPDA as well as JAs in the WT and *zmaoc2* mutant plants; however, it was slightly induced only by exogenous OPDA in the *opr3* mutant (Supplemental Figure S9e). These findings suggest that OPDA regulation of gene expression is distinct from that of JA.

## Discussion

### *ZmEREB57* participates in multiple abiotic stress response pathways in maize

Plant ERF TFs have been identified to be involved in the adaptive responses of plants to environmental stresses. Overexpression of *GmERF4* or *GmERF6* in tobacco or Arabidopsis has been demonstrated to increase tolerance to salt and drought stress (Zhang et al., 2010; Zhai et al., 2013a), and *AtABR1*, *AtRAP2.6L* and *AtRAP2.6*, which belong to the B-4 subgroup of ERFs, have been reported to respond to various biotic and abiotic types of stress (Asahina et al., 2011; Choi & Hwang, 2011; Ali et al., 2013). In the present study, ZmEREB57, a newly identified member of the ERF subfamily in maize, exhibited high sequence similarity to the Arabidopsis AtORA47 protein (Supplemental Figure S1), and its expression was induced not only by drought and salt stress, but also by ABA, JA and H_2_O_2_ treatments (Supplemental Figure S2). A *ZmEREB57* mutation in maize resulted in sensitivity to salt stress (Figure 2), whereas overexpression of *ZmEREB57* in Arabidopsis plants led to enhanced salt tolerance compared with the WT (Supplemental Figure S4). It was therefore deduced that abiotic stress induces the expression of *ZmEREB57*, and that this gene serves a role in increasing salinity tolerance.

### *ZmEREB57* regulates OPDA and JA accumulation by binding to O-box-like elements

ZmEREB57 was found to be localized to the nucleus and to exhibit transactivation activity (Figure 1), indicating that it acts as a TF, activating the expression of certain genes. It has previously been reported that AtORA47, a protein with high similarity to ZmEREB57, was able to bind to O-box elements of promoter regions (Chen et al., 2016). In the present study, we demonstrated that ZmEREB57 preferentially binds to an O-box-like motif in the promoter of the *ZmAOC2* gene, by means of Y1H, luciferase assays and EMSA (Figure 4a–c). Accordingly, a ChIP-qPCR analysis indicated that ZmEREB57 binds to O-box-like motifs to regulate the expression of the *AtAOC2* gene (Figure 4f). Furthermore, ZmEREB57 was observed to promote *ZmAOC2* expression in WT plants under conditions of salt stress (Figure 4d). AOC, a dimeric enzyme localized in chloroplasts, has been reported to be a key factor in OPDA and JA biosynthesis (Wu et al., 2011). Overexpression of *AOC* in Arabidopsis or wheat results in increased JA levels and enhanced tolerance to salt stress, indicating that this enzyme may play a role in the response to certain types of stress by regulating OPDA or JA content (Zhao et al., 2014). In our study, ZmAOC2 was confirmed to be localized in chloroplasts, and its expression had an impact on OPDA and JA levels under salt stress (Figure 5a–c), and a mutation in *ZmEREB57* resulted in decreased production of OPDA and JA levels in times of salt stress (Figure 5d and e). ZmEREB57 thus targets *ZmAOC2*, which is involved in OPDA biosynthesis, promoting OPDA and JA accumulation. Together, our data suggest that OPDA and JA biosynthesis and accumulation in maize in response to salt stress are regulated by *ZmEREB57*.

### OPDA signaling functions independently of JA signaling in the response to salt stress

JA is involved in regulating a number of plant biological processes, including development and stress responses. JA, its precursor OPDA, and associated metabolites, including MeJA and JA-Ile (collectively referred to as JAs), are all involved in mediating the response to stress caused by biotic as well as abiotic stimuli. JA and OPDA have been shown to play an important role in the adaptive response mechanisms of plants to environmental stresses (Jung et al., 2007). It is also well established that exposure to JA can improve tolerance to salinity in a number of plant species (Walia et al., 2007; Ismail et al., 2012). Furthermore, plants producing higher levels of OPDA have exhibited enhanced tolerance to drought and reduced stomatal aperture (Savchenko et al., 2014). As discussed, we deduced that ZmEREB57 promoted the accumulation of OPDA and JA by regulating *ZmAOC2* expression, and OPDA and JA content was observed to be lower in *zmereb57* mutants than in WT plants under conditions of salt stress. The observation that exogenous application of OPDA and JA to the plants could restore the phenotype of *zmereb57* mutants to one more tolerant of salt stress (Figure 6), indicates that OPDA and JA play a positive role in the response of maize to high salinity conditions.

The expression of genes regulated by JAs is altered in response to different types of stress, and COI1 is considered to be a central component of signaling pathways involving JAs (Taki et al., 2005). OPDA also induces a change in expression of certain genes, and transcription analyses have revealed distinct sets of genes that are regulated specifically by OPDA, but not JA (Stintzi et al., 2001). We used RNA-seq analysis to identify genes that respond to OPDA or JA treatment (Figure 7). The DEGs shared between JA and OPDA stimulation include signaling components, genes involved in metabolic pathways, and TFs, whereas the characteristics of J&NRGs and O&NRGs differ greatly (Figure 6 and Supplemental Figure S7). Treatment of plants with exogenous OPDA or JA was observed to restore the phenotype of strains conferring mutations in *AOC2* or *OPR3* under salt stress conditions (Figure 8). Saline treatment led to an increase in the expression of OPDA-specific response genes in *zmopr3* mutants, but not *zmaoc2* mutants, while addition of exogenous OPDA did induce their expression in the *zmaoc2* mutants (Supplemental Figure S9a–e). The activation of the *AtVSP2* gene requires the presence of JA (TaKi et al., 2005), and we found that, under conditions of salt stress, levels *ZmVSP9a* transcript were lower in both mutants that were not able to accumulate JA compared with the WT plants. Only treatment with exogenous JA, and not OPDA, resulted in restoring the expression of *ZmVSP9a* to that observed in the WT, and this occurred only in the *zmopr3* mutant strains (Supplemental Figure S9f). These findings suggest that OPDA functions as a signaling mediator in the maize salt response through a mechanism that is distinct from that of JAs.

A total of 4,032 genes were differentially expressed as a result of salt stress as well as OPDA treatment, including a number of signal transduction components and TFs (Figure 6 and Supplemental Figure S7), suggesting that OPDA and its associated response genes are required in the response mechanism to saline conditions. For example, ABA regulates ion channel activity and the stomatal opening of plant guard cells under conditions of drought or salt stress, and serves a role in the transcriptional and post-transcriptional regulation of certain stress-response genes (Wu et al., 2009). Furthermore, many TFs, such as those belonging to the bZIP, MYB, NAC and WRKY families, may function as activating or inhibiting factors participating in salt stress responses through an ABA-mediated pathway (Jakoby et al., 2002; Hiroshi et al., 2003; Marè et al., 2004; Thoenes et al., 2004; Hu et al., 2010). It is commonly known that environmental factors, such as drought, heat, cold and salinity, cause oxidative stress in plants (Xiong et al., 2002) and mitogen-activated protein kinase is a critical regulator of the antioxidant defense system in response to such stimuli, as well as ABA, brassinosteroids and reactive oxygen species (Chardin et al., 2017). The expression of many of the OPDA-specific response genes identified in this study is regulated in response to salt stress. These results suggest that OPDA modulates a distinct set of genes in response to stress via independent as well as common pathways to JA signaling.

Based on the results, a functional model of ZmEREB57 response to salt stress was proposed (Figure 9). ZmEREB57 may have a unique value in its ability to confer salt tolerance to *Z. mays* and other plant species because of its considerable upregulation in response to salt. ZmEREB57 serves a transcription factor that binds to *cis*-acting motifs O-box in the promoter to regulate the transcript of the genes, including *ZmAOC2*, etc. The induction of *ZmAOC2* significantly increased the biosynthesis of OPDA and JA. OPDA and JA mediate regulation of the salt-responsive genes through COI-independent and COI1-dependent manner, respectively. In this way, it conferred increased tolerance to salt stress in *Z. mays*.

**Figure 9.**
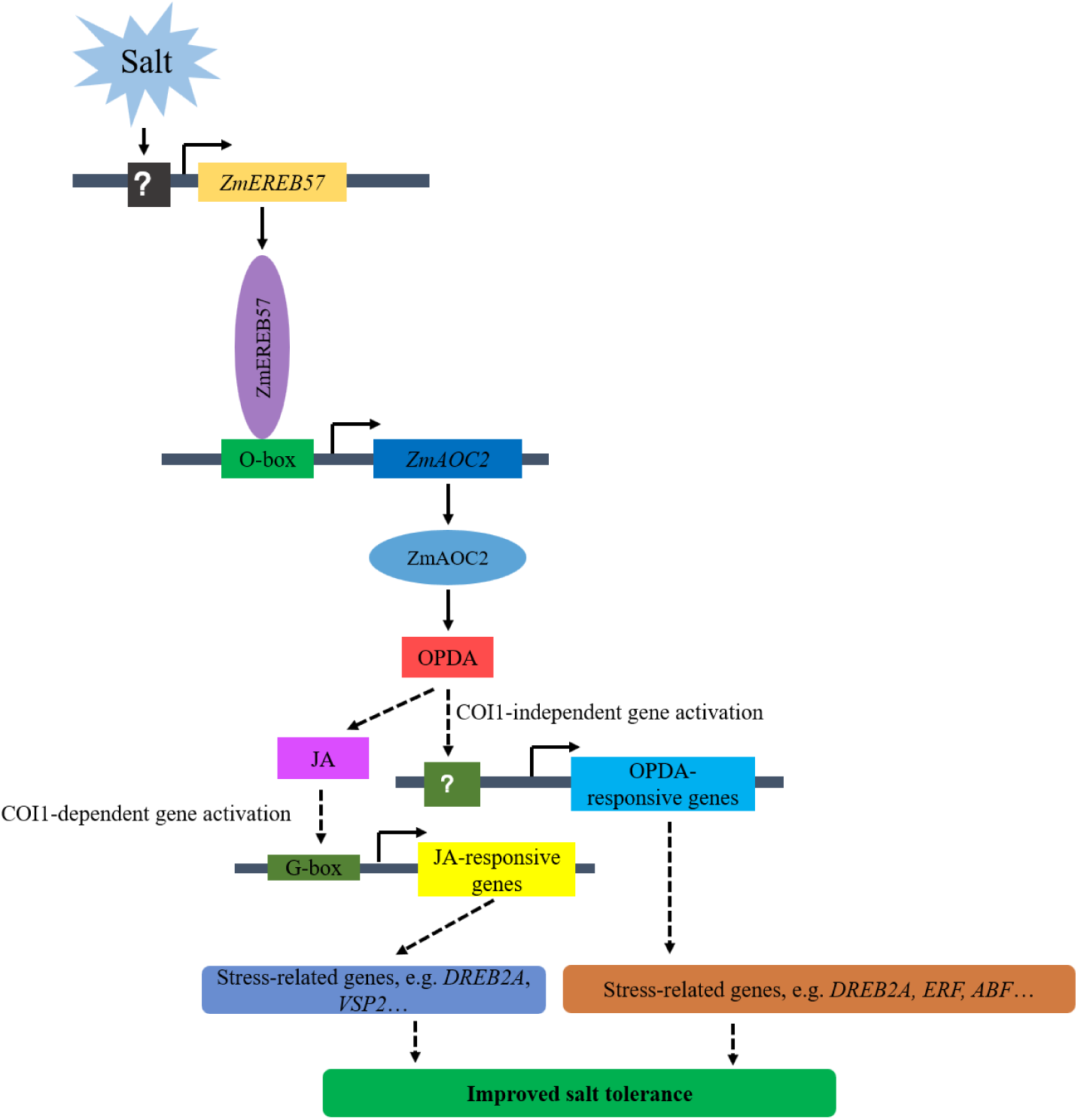
A proposed model describing the regulation mechanism of *ZmEREB57* in response to salt stress. Salt stress induces the expression of *ZmEREB57*, and ZmEREB57 activates the transcription of *ZmAOC2* by directly binding to O-box element in the promoter. The induction of *ZmAOC2* leads to increased OPDA and JA accumulation, and OPDA directly via a COI1-independent or indirectly JA-COI1-independent path regulates the expression of stress-related genes, which confer increased tolerance to salt stress in *Z. mays*.

## Materials and methods

### ZmEREB57 phylogenetic analysis

*ZmEREB57* full-length cDNA was amplified by PCR with gene-specific primers listed in Table S8. Total five sequences, including the full open reading frame of *ZmEREB57* and homologous genes in maize (*Zea mays*), rice (*Oryza sativa*), Arabidopsis (*Arabidopsis thaliana*), *Brachypodium distachyon*, and *Glycine Max*, were obtained from NCBI (https://www.ncbi.nlm.nih.gov/). The resulting encoded amino acid sequences were aligned in ClustalW (2.0) (http://www.clustal.org/) and a phylogenetic tree was constructed in MEGA v5.1 (https://megasoftware.net/), using the neighbor-joining with 1000 bootstrap replicates in p-distance model method.

### Subcellular localization and transcription activation assay

The *ZmEREB57* or *ZmAOC2* coding region was cloned into the pBI121-GFP vector by means of a Gateway LR reaction (Invitrogen, the gene-specific primers listed in Supplemental Table S8) to produce the *35S*:*ZmEREB57*-GFP or *35S*:*ZmAOC2*-GFP construct. The empty *35S*:GFP vector was used as a control. The transient expression of the green fluorescent protein (GFP)-fused proteins in tobacco (*Nicotiana tabacum)* epidermal cells was performed as described by Yoo et al. (2007) and transfected cells were observed using a confocal laser scanning microscope (Leica TCS SP2; Leica Microsystems GmbH).

For transcription activation analysis, full-length *ZmEREB57* was ligated into the *EcoRI* and *BamHI* sites of the GAL4 DNA-binding domain (BD) expression vector (Clontech) by the gene-specific primers (Supplemental Table S8) and transformed into AH109 yeast cells with pGADT7. The interaction between pGBKT7-53 and pGADT7-LargeT was used as a positive control, and the empty BD vector was used as a negative control. Positive clones were screened on the selective medium on SD/-Trp and SD/-Trp/-Leu/-Ade media. The plates were then incubated for 3 days at 28 °C.

### *ZmEREB57* expression analysis

Seeds of the maize inbred KN5585 line were surface-sterilized and germinated on filter paper for 4 days at 28°C in the dark. Seedlings with a 2-cm primary root length were transferred to Hoagland’s nutrient solution and grown in a greenhouse (14 h/10 h of light/dark) at 32°C in the day and 25°C at night until the plants reached the three-leaf stage. The seedlings were then watered with either 20% (w/v) PEG 6000, 200 mM NaCl, or 1.5 mM H_2_O_2_ solution. For phytohormone treatment, 0.1 mM ABA or 100 μM SA were added to the culture solution. The treated seedlings were harvested at 0, 3, 6, 9, 12, and 24 h, frozen immediately in liquid nitrogen, and stored at −80°C. Total RNA was extracted with TRIzol reagent (Tiangen, Beijing) according to the manufacturer’s instructions. cDNA synthesis was performed using the M-MLV reverse transcriptase (Takara Bio, Inc.) according to the manufacturer’s protocol. Quantitative (q)PCR was performed on a Bio-Rad CFX96 using the SsoFast EvaGreen Supermix (Bio-Rad), and maize Tub (NP_0011054557) was as an internal control. The sequences of the primers used in the present study are listed in Supplemental Table S8.

### Generation and analysis of maize mutant lines

Knockout Z*mereb57* mutant lines were generated on a KN5585 cultivar background, using a CRISPR-Cas9 system (Xing et al., 2014). In brief, a pCAMBIA-derived CRISPR-Cas9 binary vector with two gRNA expression cassettes targeting two adjacent sites of Z*mEREB57* was generated and transformed into *Agrobacterium* EHA105 strain, and subsequently into immature embryos of inbred KN5585 lines. To identify the positive CRISPR-Cas9 knockout lines, PCR amplicons encompassing the gRNA-targeted sites for each of the transgenic plants were sequenced by Sanger sequencing. Primers used for gene editing and plasmid construction are listed in Supplemental Table S8.

The *ZmAOC2* and *ZmOPR3* allele was isolated from the UniformMu population where the Mu-active lines were introgressed into inbred B73 genetic background (Liang et al., 2019).

### Generation of transgenic *ZmEREB57* overexpression lines in Arabidopsis

The CDS of *ZmEREB57* was inserted downstream of the CaMV *35S* promoter in the pcambia1300 vector by means of a Gateway LR reaction (Invitrogen, the gene-specific primers listed in Supplemental Table S8). The recombinant construct was infiltrated into Arabidopsis Col-0 cells by agrobacterium-mediated transformation (Clough & Bent, 1998). The antibiotic-resistant plants were selected by screening successive generations with 50 mg/L kanamycin. Two homozygous lines (T3) were chosen for the phenotypic analysis. The seeds of *atora47* (At1g74930, SALK_084382) were obtained from TAIR10. Additionally, the complementation lines of *ZmEREB57* in *atora47* mutant was also generated by overexpressing the recombinant construct.

### Phenotypic response to salt stress of transgenic and non-transgenic maize or Arabidopsis lines

To evaluate the salt stress tolerance, first the seeds of the wild-type (WT) and T3 transgenic lines were grown to the three-leaf stage in soil for three weeks, and were then adequately irrigated with a NaCl solution (200 mM) at 4 d intervals three times. Additionally, seedlings at the three-leaf stage were cultured in a hydroponic system for 6 days with Hoagland’s nutrient solution supplemented with 200 mm NaCl solution. The nutrient solutions were aerated with a mini air pump and supplemented with fresh solution to maintain volume. For the phytohormone treatments, 100 μM OPDA and jasmonic acid were added to the culture solution.

Transgenic and non-transgenic Arabidopsis seeds were surface sterilized by immersion in 0.1% (w/v) mercuric chloride and grown on Murashige and Skoog (MS) solid medium. The plates were kept at 4°C in the dark for 3 days and then transferred to conditions of a 16-h photoperiod (light intensity, 200 mM m^−2^ s^−1^) at 22°C and 70% relative humidity. Four-day-old seedlings were planted on fresh MS solid medium containing 200 mM NaCl and grown for 10 days. The plants were then examined for their phenotypic characteristics, leaf water potential and chlorophyll content as described previously (Ni et al., 2019).

### DAP-seq assay

The DNA affinity purification sequencing (DAP-seq) experiments and data analysis were conducted as previously described with minor modification (Bartlett et al., 2017). First, genomic DNA (gDNA) was extracted from young leaf tissue of maize. Then a gDNA DAP-seq library was prepared by attaching a short DNA sequencing adaptor to the purified and fragmented gDNA. The adapter sequences were truncated Illumina TruSeq adapters; the TruSeq Universal and Index adapters corresponded to the DAP-Seq Adapter A, and Adapter B. The DAP-seq gDNA library was prepared using KAPA HiFi HotStart ReadyMix with unique index primers. ZmEREB57 was fused to HaloTag using the kit from pFN19K HaloTag T7 SP6 Flexi Vecto (cat. no. G184A) (Promega). ZmEREB57 fused to HaloTag was expressed using the TnT SP6 High-Yield Wheat Germ Protein Expression System (L3260) (Promega), and then purified using Magne HaloTag Beads (G7281) (Promega). The Magne HaloTag Beads and ZmEREB57-HaloTag mixture were incubated with 500 ng DNA library in 40 µL PBS (Phosphate Buffered Saline) buffer with slow rotation in a cold room for 1.5 h. The beads were washed five times with 200 µL PBS + NP40 (0.005%), resuspended in PBS buffer, the supernatant was removed, and 25 µL EB buffer was added and samples were incubated for 10 min at 98 ◦C to elute the bound DNA from the beads. The correct DAP-seq library concentration to achieve a specific read count was calculated on the basis of on library fragment size. Negative-control mock DAP-seq libraries were prepared as described above, without the addition of protein to the beads. The final products were selected from 250 to 500 bp for ChIP-seq sequencing. We defined target genes as those that contained DAP-seq peaks located within the transcribed regions of genes, introns, or 2 kb upstream from the transcription start site (TSS), or 2 kb downstream from the transcription termination site (TTS). DAP-seq reads were aligned to the maize genome using Bowtie 2 (Langmead & Salzberg, 2012), which supports gapped and paired-end alignment modes. We ran Bowtie version 2.2.3 with default parameters and reported only unique alignments. DAP-seq peaks were detected by MACS2 (Zhang et al., 2008). We used MACS version 2.0.10 with default parameters, as duplicates were allowed, and the *q*-value < 0.05. Core motifs were identified by MEME-ChIP (Machanick & Bailey, 2011).

### Yeast one-hybrid (Y1H) assay

The coding sequence of *ZmEREB57* was obtained and digested by *EcoRI* and *BamHI* and inserted into the pGADT7-AD vector (Clontech) containing the GAL4 active domain. The promoters of *ZmAOC2* were cloned into the *KpnI* and *XhoI* sites of the pHiS2.1 vector. Plasmids were transformed in pairs in the yeast AH109 strain, which was then selected on SD/-Trp/-Leu and SD/-Trp/-Leu/-His media. The plates were then incubated for 3 days at 28°C. All the primers used for constructs are listed in Supplemental Table S8.

### Luciferase assay

Full-length *ZmEREB57* and promoter fragments of *ZmAOC2* were inserted into the 35S:GFP and pGreenII 0800:LUC vectors (the gene-specific primers listed in Supplemental Table S8), respectively. The recombinant vectors were transformed into tobacco (*Nicotiana benthamiana*) leaves using the agrobacterial infiltration method (*A. tumefaciens strain* GV3101). The luciferase signal was detected at 72 h post-transfection using a Tanon 5200 multi-chemiluminescence imaging system (Tanon Science & Technology Co.).

### Electrophoretic Mobility Shift Assay (EMSA)

The full-length *ZmEREB57* coding region was inserted into the pET30a vector for expression. The recombinant fusion plasmid was transformed into the *Escherichia coli* BL21(DE3) strain and overexpression of the cloned genes was induced with 0.5 mM isopropyl-β-D-thiogalactoside at 37°C for 7 h. For isolation and purification of the recombinant protein, bacterial cells were pelleted after induction, resuspended in 10 mL ice-cold binding buffer (0.5 M NaCl, 20 mM Tris-HCl, 5 mM imidazole, pH 7.9), and sonicated on ice for 10 min (30 s pulse/min), until the samples were no longer viscous. Following centrifugation at 12,000 × g for 15 min at 4°C, supernatants were collected, loaded onto His-Bind^®^ Resin columns (EMD Millipore; Merck KGaA), and the recombinant ZmEREB57 protein was eluted with elution buffer (0.5 M NaCl, 20 mM Tris-HCl, 1 M imidazole, pH 7.9). The 5’-end biotin probes were generated using a DIG Gel Shift Kit (Roche, China) (Supplemental Table S8), and the probes without a biotin label were set as a competitor probe. EMSAs for detecting interactions between ZmEREB57 and *ZmAOC2* promoter were performed as described by Liu et al. (2006).

### Chromatin immunoprecipitation (ChIP)-qPCR assay

ChIP assays were performed as reported previously (Xiang et al., 2021). After two weeks growth period, 1.5g of transgenic *ZmEREB57*-overexpressing or WT Arabidopsis plants were harvested after treated with or without 100 mM NaCl for 7 d, and fixed in 1% formaldehyde for 10 min with vacuum infiltration. Crosslinking was quenched with the addition of glycine to a final concentration of 0.125 M. Chromatin was sheared to 200 to 500 bp by ultrasonic treatment (Bioruptor; Picoruptor; 15 cycles of 30 s on and 30 s off) and immunoprecipitated using an anti-GFP antibody (Sigma-Aldrich; Merck KGaA). The immunoprecipitated and purified DNA was used for quantitative real-time PCR, and all the samples were diluted to 10 ng μL^−1^ and reacted with 5 μL of SYBR Premix Ex Taq (2×), 0.2 μL of PCR forward primer (10 μm), 0.2 μL of PCR reverse primer (10 μm), and 1 μL of DNA template. The primers used to amplify the enriched region of the target genes are listed in Supplemental Table S8. A fragment of the *Actin* coding region was used as the internal reference gene. Determined values of quantity were normalized with the amount of input DNA.

### Detection of phytohormones

Phytohormones contents were detected by MetWare (http://www.metware.cn/) based on the AB Sciex QTRAP 6500 LC-MS/MS platform. For each replicate, approximately 2 g (fresh weight) of fresh maize samples, including KN5585 and mutants, were harvested at three-leaf stage after treated with or without 200 mM NaCl, immediately frozen and homogenized under liquid nitrogen, and stored at −80°C until further use. Each sample was dissolved in 3 ml of, 2 propanol/H2O/concentrated HCl (2:1:0.002, vol/vol/vol) extraction solvent, and H_2_JA (TCI AMERICA, Portland, OR, USA) was added as an internal standard. The mixture was vortexed for 10 min and centrifuged at 13,000 g at 4°C for 5 min. The lower phase was transferred to clean plastic microtubes. The samples were allowed to evaporate to dryness, dissolved in 100 μL 80% methanol (v/v), and filtered through a 0.22-μm membrane filter for liquid chromatography-mass spectrometry (LC-MS/MS) analysis. A linear ion trap orbitrap mass spectrometer (Orbitrap Elite; Thermo Fisher Scientific, Waltham, MA, USA) coupled online with an UHPLC system (ACQUITY UPLC; Waters) was used to perform the quantitation. JA and OPDA were separated using a C18 column (Waters, Milford, MA, USA) operated in the negative ion mode (Pan et al. 2010). OPDA (Item No. 88520, Cayman Chemical, Ann Arbor, Michigan, USA) and JA (Product n o.14631, Sigma Aldrich, Gillingham, UK) were used as the standards. A statistical analysis. Elkhart, IN, USA). Data were determined from three biological replicates.

### RNA-seq analysis

The KN5585 WT plants were treated with or without 200 mM NaCl or OPDA or JA at the three-leaf stage. Approximately 0.5 g leaves (fresh weight) were collected and immediately frozen in liquid nitrogen, with three biological replicates taken for each treatment. Total RNA was extracted using TRIzol reagent (TIANGEN BIOTECH, Beijing, China). and the cDNA library was constructed with the NEBNext Ultra RNA Library Prep Kilt for Illumina (NEB, Ipswich, MA, USA), according to manufacturer’s instructions. Cutadapt software (https://cutadapt.readthedocs.io/) was used to exclude reads that contained adapter contamination, as well as low-quality and undetermined bases. RNA-seq data were normalized by rounding fragments per kilobase million values. The differentially expressed mRNAs were identified by selecting those with fold-changes >2 or <0.5 and p-value<0.05, using packages edgeR (https://bioconductor.org/packages/release/bioc/html/edgeR.html) or DESeq2 (https://bioconductor.org/packages/release/bioc/html/DESeq2.html) in R (https://www.r-project.org/) (Anders and Huber, 2010), followed by gene ontology (GO) term enrichment analysis (Young et al., 2010).

### Statistical analysis

A two-way ANOVA and Tukey’s post hoc test were used to compare measurements and characteristics of plant genotypes tested in the present study. A *p*-value of <0.05 was considered to confer a significant difference.

## Acknowledgments

We thank Fanguo Chen (Shandong University) for reviewing the manuscript and providing excellent suggestions for improvement. We thank the ChinaMu Project for the maize mutants.

## Authors’ contributions

J.Z and H.L designed the study. J.Z and X.W performed most of experiments. J.Z wrote up the manuscript. J.Y and H.L made fine revision on the manuscript. C.Y, H.Z, and J.Y performed a part of the experiment and analyzed the data. All authors read and approved the manuscript.

## Supplementary materials

**Supplemental Figure 1.** Sequence analysis of ZmEREB57 compared with related AP2/ERF proteins. (a) Amino acid sequence alignment encompassing the conserved AP2/ERF domain. The locations of the three β-sheets and one α-helix of the AP2/ERF domain are indicated above the sequences. (b) Neighbor-joining phylogenetic tree of AP2/ERF transcription factors, including ZmEREB57, built using the amino acid sequence alignment. (c) Predicted three-dimensional structures of the of ZmEREB57 and AtORA47 AP2/ERF domains. AP2/ERF, APETALA2/Ethylene-Responsive Factor; AtORA47, *Arabidopsis thaliana* octadecanoid-responsive AP2/ERF-domain TF 47; ZmEREB57, *Zea mays* Ethylene Responsive Element Binding Factor 57.

**Supplemental Figure 2.** Expression analysis of *ZmEREB57* under stress and hormone treatments. Seedlings at the three-leaf stage were treated with different types of abiotic stress and phytohormones, and *ZmEREB57*expression levels in the leaves were determined by reverse transcription-quantitative PCR. Bars represent mean ± SD (n = 3 replicates). ABA, abscisic acid; JA, jasmonate; PEG, polyethylene glycol; ZmEREB57, *Zea mays* Ethylene Responsive Element Binding Factor 57.

**Supplemental Figure 3.** Expression analysis of *ZmEREB57* in WT maize and *zmereb57* mutants. ZmEREB57 expression levels in WT maize and *zmereb57* mutants were determined by reverse transcription-quantitative PCR(a) and PCR (b). *ZmTUB* served as a control. Different letters indicate values which departed significantly from those of WT maize.

**Supplemental Figure 4.** *ZmEREB57*-overexpressing transgenic Arabidopsis lines and their response to salt stress. (a) Phenotype of seedlings of WT and transgenic lines that were germinated and transferred to MS medium supplemented with or without 100 mM NaCl. (b) Root length and (c) fresh weight of WT and transgenic lines exposed to 0 and 100 mM NaCl treatment. (d) Phenotype of WT and transgenic plants before (Control) and after 200 mM NaCl treatment for 7 days. (e) Fresh weight and (f) chlorophyl content of WT and transgenic lines before (Control) and after 200 mM NaCl treatment for 7 days. Results are presented as mean values of three replicates ± SD (n > 6 each). Scale bar = 1 cm. Different letters indicate values which departed significantly from those of WT maize. AtORA47, *Arabidopsis thaliana* octadecanoid-responsive AP2/ERF-domain TF 47; WT, wild type; ZmEREB57, *Zea mays* Ethylene Responsive Element Binding Factor 57.

**Supplemental Figure 5.** ZmEREB57 binds directly to the *AtAOC2* promoter. (a) Luciferase assay for the *AtAOC2* promoter. The pro *AtAOC2*:LUC-35S:REN reporter construct was transiently expressed in tobacco leaves together with the control vector or the effector, and images of luciferase signal detection were captured. (b) An Electrophoretic Mobility Shift Assay indicates that ZmEREB57 binds to the *AtAOC2* promoter *in vitro*. His-ZmEREB57 protein (2 μg) was incubated with labeled probe. Unlabeled (competitive) probe at 25x and 50x molar excess competed with the labeled probe.

**Supplemental Figure 6.** Maize *ZmEREB57* knockout alleles exhibit increased sensitivity to salinity. (a) Appearance of 2-week-old *zmereb57*-1, *zmereb57*-2 and WT plants grown in Hoagland’s nutrient solution before and after treatment with 200 mM NaCl at 48 h. (b) Fresh weight, (c) chlorophyll content, (d) OPDA levels and (e) JA levels in the three strains under control or salt-treatment conditions. Results are presented as the mean ± SD (n = 3 replicates). Different letters indicate values which departed significantly from those of WT maize. JA, jasmonate; OPDA, 12-oxo-phytodienoic acid; WT, wild type; ZmEREB57, *Zea mays* Ethylene Responsive Element Binding Factor 57.

**Supplemental Figure 7.** Global transcriptome responses of maize to salt stress alone or in combination with phytohormone treatment. (a) Gene ontology term enrichment annotation and (b) Kyoto Encyclopedia of Genes and Genomes pathway enrichment annotation of the 3, 686 DEGs identified to be shared among the three types of treatment: 200 mM NaCl supplemented with OPDA and 200 mM NaCl supplemented with JA. (c) Gene ontology term enrichment annotation and (d) Kyoto Encyclopedia of Genes and Genomes pathway enrichment annotation of the 2, 681 DEGs identified to be shared among the three types of treatment: 200 mM NaCl alone, 200 mM NaCl supplemented with OPDA, and 200 mM NaCl supplemented with JA. Numbers next to each bar represent the number of DEGs. DEG, differentially expressed gene; JA, jasmonate; OPDA, 12-oxo-phytodienoic acid.

**Supplemental Figure 8.** Relative expression levels of stress-response genes in WT and *zmereb57* mutants under conditions of salt stress. Results are represented as the mean ± SEM (n = 3 seedlings). Different letters indicate values which departed significantly from those of WT maize. WT, wild type; ZmEREB57, *Zea mays* Ethylene Responsive Element Binding Factor 57.\

**Supplemental Figure 9.** Relative expression of OPDA-specific response genes in maize WT plants and mutant strains deficient in OPDA and JA biosynthesis genes under different treatments. Results are represented as the mean ± SEM (n = 3 seedlings). Different letters indicate values which departed significantly from those of WT maize. JA, jasmonate; OPDA, 12-oxo-phytodienoic acid; WT, wild type.

**Supplemental Table 1** The relative abundance of DEGs induced by 200 mM NaCl treatment.

**Supplemental Table 2** Identification and analysis of ZmEREB57 direct target genes.

**Supplemental Table 3** The relative abundance of DEGs induced by 100 μM JA treatment.\

**Supplemental Table 4** The relative abundance of DEGs induced by 100 μM OPDA treatment.

**Supplemental Table 5** The relative abundance of common DEGs uniquely induced by NaCl & OPDA treatment.

**Supplemental Table 6** The relative abundance of common DEGs uniquely induced by NaCl & JA treatment.

**Supplemental Table 7** The relative abundance of common DEGs uniquely induced by NaCl & OPDA & JA treatment.

**Supplemental Table 8** Primers and their sequences used in this study.

